# Tuning antiviral CD8 T-cell response via proline-altered peptide ligand vaccination

**DOI:** 10.1101/862144

**Authors:** Adil Doganay Duru, Renhua Sun, Eva B. Allerbring, Jesseka Chadderton, Nadir Kadri, Xiao Han, Hannes Uchtenhagen, Chaithanya Madhurantakam, Sara Pellegrino, Tatyana Sandalova, Per-Åke Nygren, Stephen J. Turner, Adnane Achour

**Affiliations:** Science for Life Laboratory, Department of Medicine Solna, Karolinska Institute, and Division of Infectious Diseases, Karolinska University Hospital, Solna, Stockholm, Sweden; NSU Cell Therapy Institute & Dr. Kiran C. Patel College of Allopathic Medicine, Nova Southeastern University, Fort Lauderdale, FL, USA; Department of Microbiology, Biomedical Discovery Institute, Monash University, Clayton, Australia; DISFARM, Dipartimento di Scienze Farmaceutiche, Sezinone Chimica Generale e Organica, Università degli Studi, Milano, Italy; Division of Protein Engineering, Department of Protein Science, School of Engineering Sciences in Chemistry, Biotechnology and Health, AlbaNova University Center, Royal Institute of Technology, Stockholm, Sweden

**Keywords:** LCMV, CD8 T cells, Altered Peptide Ligands, TCR, MHC class I, vaccination, immune escape, X-ray crystallography

## Abstract

Viral escape from CD8^+^ cytotoxic T lymphocyte responses correlates with disease progression and represents a significant challenge for vaccination. Here, we demonstrate that CD8^+^ T cell recognition of the naturally occurring MHC-I-restricted LCMV-associated immune escape variant Y4F is restored following vaccination with a proline-altered peptide ligand (APL). The APL increases MHC/peptide (pMHC) complex stability, rigidifies the peptide and facilitates T cell receptor (TCR) recognition through reduced entropy costs. Structural analyses of pMHC complexes before and after TCR binding, combined with biophysical analyses, revealed that although the TCR binds similarly to all complexes, the p3P modification alters the conformations of a very limited amount of specific MHC and peptide residues, facilitating efficient TCR recognition. This approach can be easily introduced in peptides restricted to other MHC alleles, and can be combined with currently available and future vaccination protocols in order to prevent viral immune escape.

**Author Summary:** Viral escape mutagenesis correlates often with disease progression and represents a major hurdle for vaccination-based therapies. Here, we have designed and developed a novel generation of altered epitopes that re-establish and enhance significantly CD8^+^ T cell recognition of a naturally occurring viral immune escape variant. Biophysical and structural analyses provide a clear understanding of the molecular mechanisms underlying this reestablished recognition. We believe that this approach can be implemented to currently available or novel vaccination approaches to efficiently restore T cell recognition of virus escape variants to control disease progression.

## Introduction

Recognition of major histocompatibility complex class I (MHC-I)-restricted viral peptides is a prerequisite for CD8^+^ T-cell activation, control and/or clearance of viral infections. Usually, cytotoxic T-lymphocyte (CTL) responses are directed towards a limited number of immunodominant viral peptides [1] and selection pressure imposed by adaptive immune responses leads often to the emergence of viral populations with a limited number of recurring escape mutations [2–4]. Epitope mutations can impair CTL responses [5] by *e.g*. altering antigen processing [6, 7], reducing the overall stability of peptide/MHC complexes (pMHC) [8, 9] and/or disrupting T-cell receptor (TCR) recognition [10, 11]. CTL escape variants correlate with disease progression [12, 13] and represent a major hurdle for disease control as well as for the design of T-cell based vaccines [14].

To our knowledge, previous use of wild-type and escape epitopes in vaccination experiments has not provided efficient CTL responses against MHC-restricted viral escape variants [14, 15]. Therefore, the design of altered peptide ligands (APLs) that could promote such responses would represent a crucial step towards the development of efficient vaccines [16]. Although our understanding of the interactions between TCRs and pMHC has increased dramatically, the impact of individual peptide modifications on TCR recognition remains difficult to predict. Even subtle peptide alterations can significantly impact on pMHC stability, and impair or abolish T cell recognition. A conventional and sometimes successful approach to design APLs with enhanced pMHC stability and immunogenicity has been to optimize interactions between peptide anchor residues and MHC binding pockets [17–19]. However, escape variants that target TCR recognition often exhibit optimal MHC anchor residues, reducing possibilities for such modifications.

Optimally, the introduced modifications should also not alter the conformation of APLs compared to the original epitopes, in order to elicit efficient cross-reactive CTL responses towards the wild-type epitope [18, 20, 21]. Therefore, the design of a novel generation of APLs that could promote such responses would represent a crucial step towards the development of efficient anti-viral T-cell based vaccines [22].

We have previously demonstrated that the immunogenicity of the cancer-associated H-2D^b^-restricted antigen gp100_25-33_ [23] or the T cell epitope associated with impaired peptide processing (TEIPP) neo-epitope Thr4 [24–26] was dramatically improved following substitution of peptide position 3 to a proline (p3P). Comparative structural analyses revealed that the conformation of the APLs was similar to wild-type epitopes, and that the stabilizing effect of p3P is accounted for by van der Waals and CH-π interactions with the H-2D^b^ residue Y159, conserved among most known mouse and human MHC-I alleles, resulting in rigidification of the pMHC complex [27]. Importantly, vaccination with p3P-modified APLs elicited high frequencies of CTLs from the endogenous repertoire that efficiently targeted H-2D^b^/gp100_25-33_ and H-2D^b^/Trh4 complexes on melanoma cells [23, 24].

In the present study, we addressed if we could restore endogenous T cell recognition of a naturally occurring viral escape variant following vaccination with a p3P-modified APL. It is well established that infection of C57/Bl6 mice with LCMV induces robust CTL responses towards the immunodominant H-2D^b^-restricted epitope gp33 (KAV**<u>Y</u>**NFATM) [28]. Upon CTL pressure, a limited number of mutations in gp33 emerge, with consistent patterns, allowing for viral CD8^+^ T-cell escape [2, 4, 29, 30]. The main naturally occurring mutation that allows LCMV to efficiently escape immune recognition, is the Y4F substitution (KAV**<u>F</u>**NFATM) which abrogates endogenous CD8^+^ T cell recognition as well as recognition by the H-2D^b^/gp33-specific TCR P14. Here, we demonstrate that peptide vaccination with PF (KA**<u>PF</u>**NFATM) restores P14 recognition of Y4F in LCMV-infected mice. Furthermore, vaccination with influenza constructs that encode for PF provokes significant endogenous CD8^+^ T cell cross-recognition of Y4F. Comparison of crystal structures of an ensemble of pMHC complexes before and after binding to the TCR P14 revealed that i) P14 binds nearly identically to all complexes, ii) the conformations of peptide residues p1K and p6F as well as H-2D^b^ residues R62, E163 and H155 are affected by the p3P modification, predisposing pMHC complexes for enhanced TCR recognition. The p3P modification also decreases the entropic penalty for TCR recognition. In conclusion, our results demonstrate the possibility to vaccinate with modified peptides and/or proteins for enhanced T cell recognition, and may form an alternative basis for novel strategies to target viral escape mutants.

## Results

### The p3P modification enhances pMHC stability without altering structural conformation, restoring P14 TCR recognition

Circular dichroism (CD) measurements revealed a consistent increase in pMHC complex thermal stability for the p3P-substituted peptides V3P (KA<u>PY</u>NFATM) and PF (KA<u>PF</u>NFATM) compared to the wildtype gp33 (KA<u>VY</u>NFATM) and escape variant Y4F (KA<u>VF</u>NFATM) epitopes, respectively (Fig. 1A, Table 1). Importantly, the H-2D^b^/gp33 and H-2D^b^/Y4F display equivalent thermal stability (Fig. 1A). Furthermore, surface plasmon resonance (SPR) analyses revealed significantly higher binding affinity of soluble P14 TCR to H-2D^b^/V3P and H-2D^b^/PF compared to H-2D^b^/gp33 and H-2D^b^/Y4F, respectively (Fig. 1B, Table 1). In contrast to an undetectable affinity to H-2D^b^/Y4F, P14 bound to H-2D^b^/PF. Interactions between soluble P14 and H-2D^b^/gp33 and H-2D^b^/V3P were also characterized using isothermal titration calorimetry (ITC), revealing that binding of P14 to H-2D^b^/V3P was mainly enthalpy-driven with almost null contribution of entropy, whereas binding to H-2D^b^/gp33 was entropically unfavorable (Fig. S1, Table 1). In conclusion, the p3P modification increases pMHC stability, resulting in recognition of PF by P14 and enhances TCR affinity by decreasing the entropic cost for binding.

**Figure 1.**
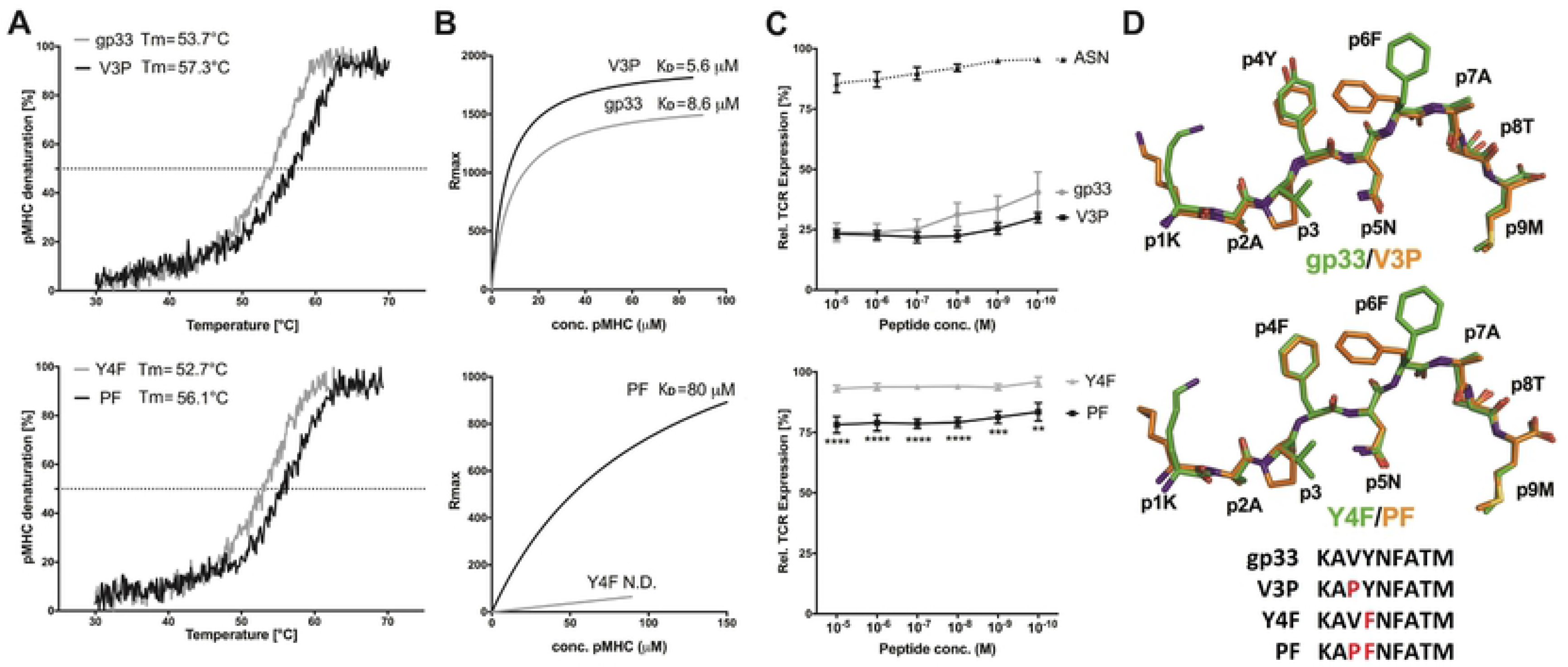
The p3P modification enhances pMHC stability without altering structural conformation, reestablishing TCR recognition. **A. The p3P modification increases pMHC stability.** CD melting curves of H-2D^b^/gp33 and H-2D^b^/V3P (upper panel), and H-2D^b^/Y4F and H-2D^b^/PF (lower panel). Melting temperatures (T_m_) corresponding to 50% protein denaturation are indicated. **B. The APL PF is recognized by the soluble TCR P14.** In contrast to Y4F, PF is recognized by P14. Binding affinity of the soluble TCR P14 to each pMHC was measured using SPR. K_D_ values are indicated. **C. The p3P modification increases TCR internalization.** TCR downregulation was measured following exposure of P14 T cells to H-2D^b^ in complex with each peptide at indicated concentrations on RMA cells. CD3^+^CD8^+^CD4^-^ and Vα2^+^ cells were gated to quantify TCR internalization and p values calculated by using two-way Anova with Turkey’s multiple comparison test. **** represents p<0.0001; *** 0.0002 and ** 0.0018. The H-2D^b^-restricted Influenza-derived peptide ASNENMETM (ASN) was used as negative control. **D. The p3P modification does not alter the conformation of the backbone of APLs compared to native counterparts.** Superposition of the crystal structures of H-2D^b^/V3P and H-2D^b^/PF with H-2D^b^/gp33 and H-2D^b^/Y4F, respectively, demonstrates that the p3P modification does not alter backbone conformations. Significant conformational changes are only observed for the side chains of peptide residues p1K and p6F following the p3P substitution.

Next, P14 TCR down-regulation was assessed upon exposure to gp33, V3P, Y4F or PF-loaded H-2D^b+^ RMA cells (Fig. 1C). While H-2D^b^/gp33 induced significant TCR down-regulation, none was detected with Y4F, even at high peptide concentrations. Exposure of P14 T cells to V3P equaled or increased TCR internalization compared to gp33. Importantly, exposure to PF significantly increased P14 TCR down-regulation compared to Y4F (Fig. 1C). The crystal structures of H-2D^b^/V3P and H-2D^b^/PF were determined to 2.5 and 2.6 Å resolution, respectively (Table S1), and compared with H-2D^b^/gp33 [3] and H-2D^b^/Y4F [2] (Fig. 1D, Fig. S2, Fig. S9). The overall structures of all pMHCs are nearly identical, and the amount of hydrogen bond and van der Waals interactions formed between H-2D^b^ and each p3P-APL was equivalent to each wild-type epitope counterpart. The backbone of the p3P-APL corresponds exactly to the wild-type peptides, indicating strict molecular mimicry (Fig. 1D). The root mean square deviation values for main chain atoms are 0.24-0.67Å^2^ and 0.20-0.24Å^2^ for the backbone of the H-2D^b^ heavy chain and the peptides, respectively. The only significant conformational differences between wild-type and p3P-APLs were side chain movements of peptide residues p1K and p6F towards the N-terminal and middle section of the peptide-binding cleft of H-2D^b^, respectively (Fig. 1D, Fig. S2).

### In contrast to Y4F, PF induces significant P14 T cell responses both *in vitro* and *in vivo*

First, we assessed the functional effects of all peptides on P14 T-cell activation by comparing intracellular TNF and IFNγ production, T cell degranulation (CD107a) as well as target cell lysis. CD8^+^ T cells, isolated from spleens of naïve or gp33-immunized P14 transgenic mice (P14-tg), were co-cultured with RMA cells pulsed with each peptide. Peptides gp33, V3P and PF induced significant production of TNF and IFNγ, as well as CD107a expression, while Y4F failed to induce any P14 T cell response (Fig. S3). Lysis of RMA cells by P14 T cells was also enhanced with PF compared to Y4F (Fig. S3). In conclusion, p3P-modification of the immune escape variant Y4F re-establishes *in vitro* recognition by P14 T cells (Fig. S3).

We thereafter assessed the *in vivo* impact of the p3P modification on LCMV-activated P14 T cells. 10^4^ P14 T-cells were adoptively transferred into C57Bl/6 mice, thereafter infected with the LCMV clone 13 (Fig. 2). Six days post-infection, P14 T-cells isolated from spleens (Fig. 2A–2B) were either stained with pMHC tetramers or re-stimulated with 10^-6^ M gp33, Y4F or PF. Tetramer staining demonstrated that a significant amount of the activated P14 T cells recognized the H-2D^b^/PF complex (Fig. 2C–2E). Furthermore, while PF- and gp33-stimulated P14 T-cells produced TNF and IFNγ, Y4F was not recognized (Fig. 2D–2E). Altogether, these results demonstrate that, in contrast to Y4F, PF is efficiently recognized by P14 T cells *in vivo*-activated by LCMV infection.

**Figure 2.**
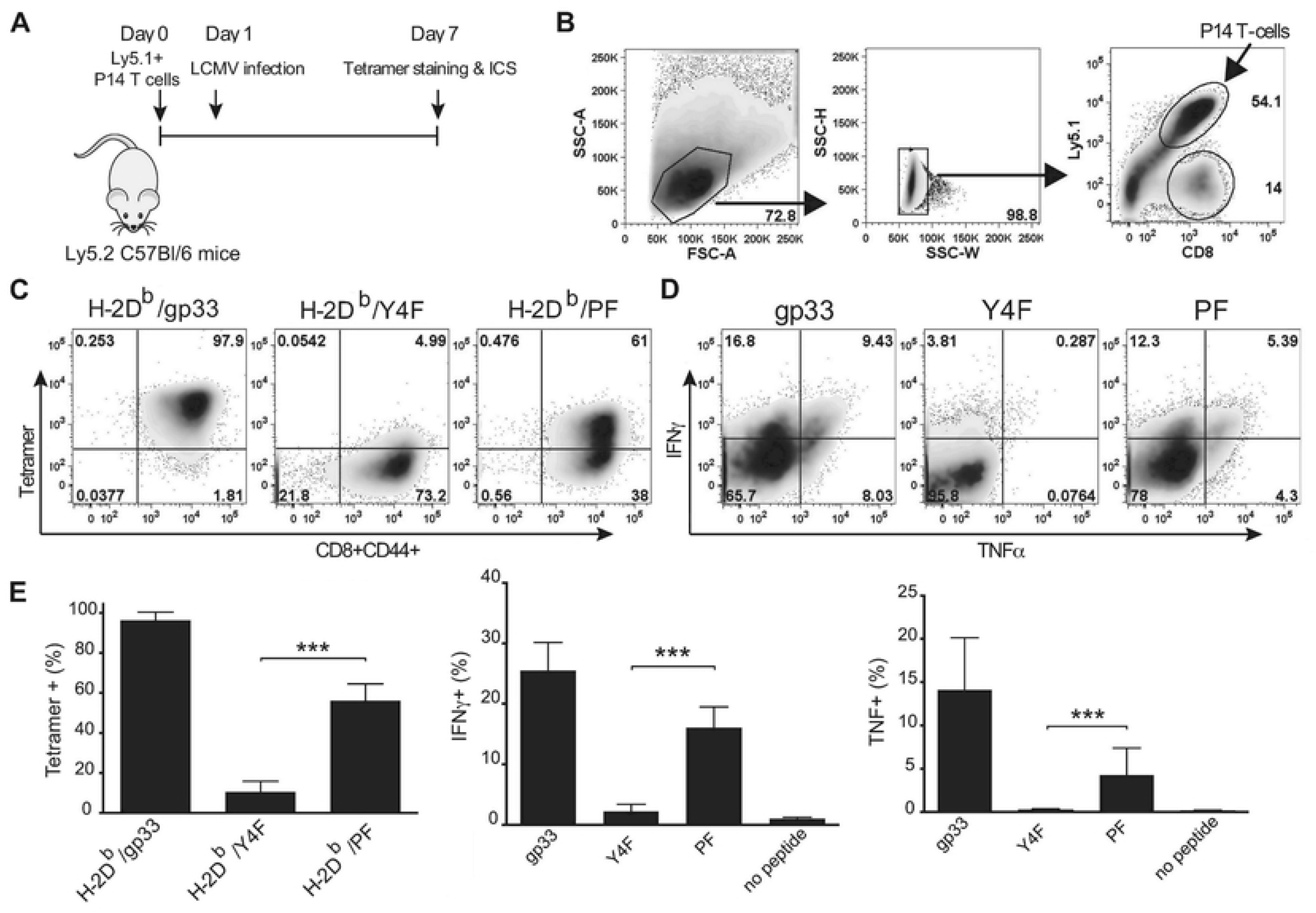
The p3P modification increases significantly P14 T cell responses. **A.** C57/Bl6 mice were adoptively transferred with 10^4^ P14 T-cells one day prior to infection with LCMV. Mice were sacrificed on day 7 post T cell transfer. T-cells from spleen were stained with PE-conjugated H-2D^b^/gp33, H-2D^b^/Y4F or H-2D^b^/PF tetramers. T cells were also stimulated with gp33, Y4F or PF peptides (10^-6^ M) for 5h, prior to assessment of intracellular IFNγ and TNF expression levels. **B.** Gating strategy used to detect CD8^+^CD44^+^ cells. P14 T cells were distinguished from endogenous T-cells using the Ly5.1 (V450) marker. **C.** Representative density plots from tetramer staining. CD8^+^ CD44^+^ P14 T-cells were stained with H-2D^b^/gp33 (left), H-2D^b^/Y4F (middle) and H-2D^b^/PF (right) tetramers. **D.** Representative ICS density plots. P14 T-cells were stimulated with peptides gp33 (left), Y4F (middle) or PF (right). **E.** CD8^+^CD44^+^ P14 T-cells from the spleen were stained with the indicated tetramers on the x-axis (left). P14 T-cells from the spleen were stimulated with the peptides indicated on the x-axis, and expression of INFg (middle) and TNF (right) was assessed. Error bars show mean +/- SD. One-way anova was performed to compare between different groups. P-values * and *** represent p<0.05 and p<0.001, respectively. The analysis was made using the GraphPad Prism software.

### Vaccination with Influenza encoding for PF activates endogenous CD8^+^ T-cells that cross-react and recognize the immune escape variant Y4F

Next, we assessed if vaccination with PF could elicit endogenous T cells that cross-react and recognize Y4F. We engineered Y4F and PF into the stalk region of the Influenza A Neuraminidase (HKx31). This well-established model results in efficient processing and presentation of epitopes on infected cells [31]. C57/Bl6 mice were infected with the modified viruses Flu(Y4F) or Flu(PF) (Fig. 3A). 10 days following infection, CD8^+^CD44^+^ splenocytes (Fig S4) were co-stained with H-2D^b^/gp33-, H-2D^b^/Y4F- and H-2D^b^/PF-tetramers (Fig. 3B–3C). Vaccination with Flu(PF) elicits endogenous T cell populations that bind to both H-2D^b^/Y4F and H-2D^b^/PF tetramers equally well (Fig. 3B). Interestingly, Flu(PF) vaccination also elicits endogenous T cell populations with dual specificity to H-2D^b^/gp33 and H-2D^b^/Y4F tetramers. In contrast, H-2D^b^/gp33-, H-2D^b^/Y4F- and H-2D^b^/PF-tetramer staining after vaccination with Flu(Y4F) failed to identify any significant T cell population (Fig. 3B).

**Figure 3.**
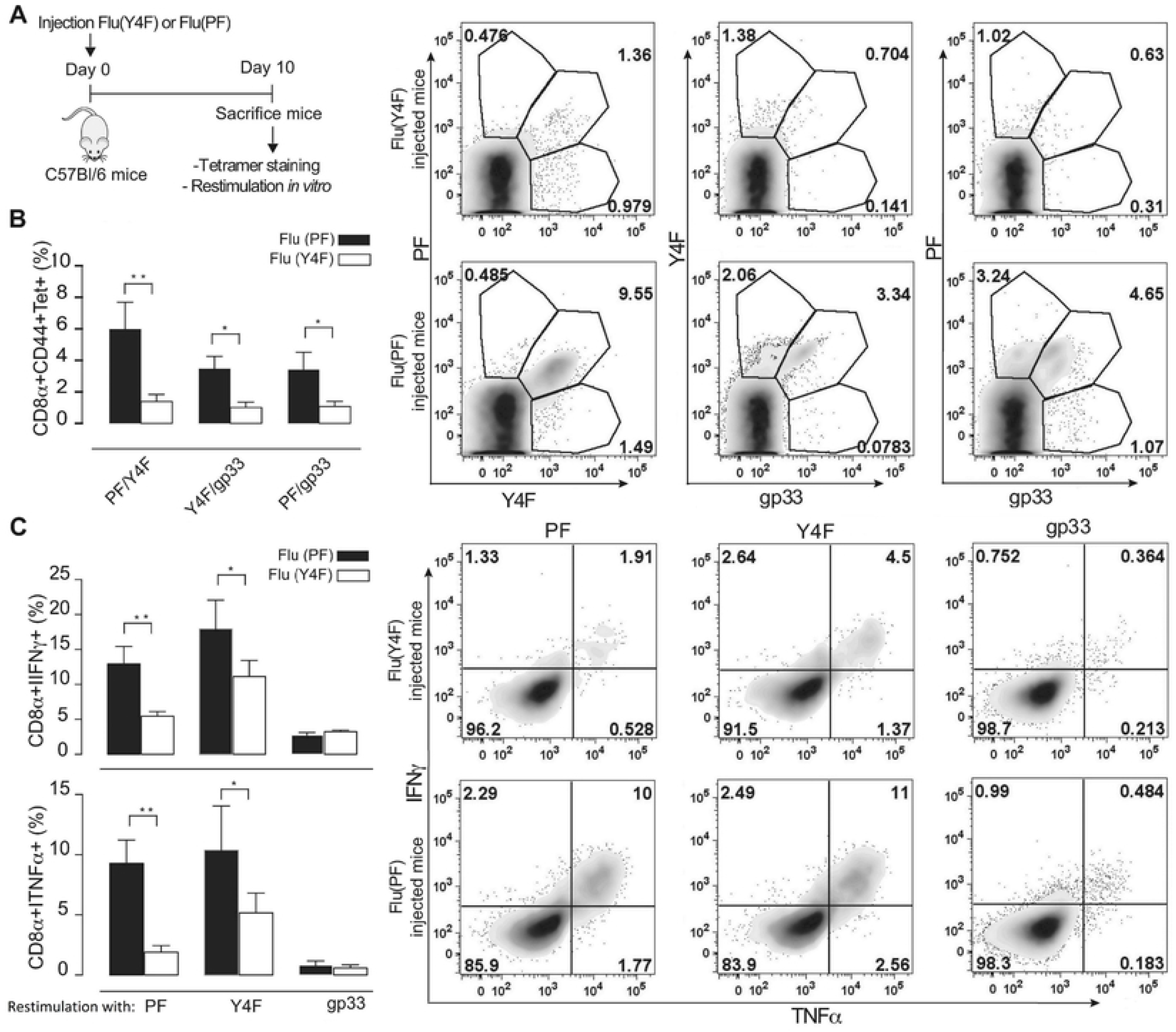
Vaccination of C57/Bl6 mice with influenza virus encoding for PF re-established efficient recognition of the immune escape variant Y4F. The escape mutant Y4F (KAV**<u>F</u>**NFATM) and the proline-modified variant PF (KA**<u>PF</u>**NF ATM) were engineered into the stalk region of neuraminidase of the Influenza A virus strain HKx31 (H3N2), and used to infect C57BL/6 mice. **A.** C57/Bl6 mice infected with either flu(Y4F) or flu(PF) were sacrificed day 10 post infection. **B.** CD8^+^ CD44^+^ cells were stained with combinations of H-2D^b^/gp33, H-2D^b^/Y4F or H-2D^b^/PF tetramers. Right top panel: Representative density plots of CD8^+^ CD44^+^ T-cells from mice infected with flu(Y4F) or flu(PF). Data from pooled 4-5 mice, representative of two different experiments. **C.** Cells were also stimulated with gp33, Y4F or PF peptides for 5 h, and intracellular IFNγ and TNF expression was determined. (Right bottom panel) CD8^+^ CD44^+^ T-cells isolated from mice infected with flu(Y4F) or flu(PF) were stimulated with either gp33, Y4F or PF (10^-6^ M), and thereafter stained for INFg and TNF. Data of IFNγ and TNF secretion from pooled 4-5 mice representative of two different experiments. Error bars show mean +/- SD. Statistical significance is presented with the p-value from a two-way Anova with Sidak’s multiple comparison test. * represents p<0.05; ** represents p<0.01. The analyses were performed using the GraphPad Prism software.

Intracellular expression of IFNγ and TNF in CD8^+^CD44^+^ endogenous T cells was assessed following stimulation with peptides gp33, Y4F or PF (Fig. 3C). In contrast to Flu(Y4F), vaccination with Flu(PF) results in significantly enhanced IFNγ and TNF levels towards both Y4F and PF. However, vaccination with neither Flu(Y4F) nor Flu(PF) induced any elicitation of IFNγ and TNF towards gp33. This is well in line with previous studies in which the Y4F-specific T cell clone YF.F3 killed efficiently targets presenting gp33 but did not produce IFNγ [32]. Similar results were obtained using bronchoalveolar lavage (BAL)-derived T cells (Fig. S4). In conclusion, vaccination with Flu(PF) induces endogenous T cell populations that respond strongly to H-2D^b^/PF and efficiently cross-react with H-2D^b^/Y4F.

### The T cell receptor P14 binds identically to H-2D^b^/gp33, H-2D^b^/V3P and H-2D^b^/PF

In order to assess the molecular bases underlying the effects of the p3P modification on T cell recognition, we determined the crystal structures of the ternary complexes P14/H-2D^b^/gp33, P14/H-2D^b^/V3P and P14/H-2D^b^/PF to 3.2, 2.8 and 1.75 Å resolution, respectively (Table S2, Fig. S5). All ternary complexes are almost identical with rmsd values of 0.5Å, 0.18-0.28Å, 0.35Å and 0.27-0.31Å for peptide clefts, peptides, TCRa and TCRb, respectively. The three ternary complexes displayed a typical TCR/pMHC binding mode with P14 diagonally positioned over the pMHC complexes (Fig. S6). The ternary structures revealed very similar TCR contacts with H-2D^b^ presenting the three different peptides, with identical conformations of the six P14 CDR loops (Fig. S6). Although CDR3a (101-**Y**G**NE**K-105) and CDR3b (93-**D**AG**GR**NTL-100) are located over the middle part of each peptide variant, only CDR3b forms hydrogen bonds with the three peptide residues p4Y, p6F and p8T (Fig. S7). All the other P14 loops CDR1a (33-E**D**S**T**F**N**-38), CDR1 b (25-NNH**D**YM-30), CDR2a (58-**L**S**V**S-61) and CDR2b (46-YS**Y**-48) interact with the H-2D^b^ heavy chain (Fig. S7).

### The immune escape mutation Y4F abrogates the hydrogen bond network formed with P14

The P14 CDR3b residues D93, G96 and R97 form a network of hydrogen bonds with the side chains of the gp33 residues p4Y and p8T, as well as with the backbone of p5N and p8T (Fig. S7). The side chain of R97β runs parallel with the peptide, stretching out to reach to the tip of p4Y, forming van der Waals interactions with the side chain of p6F, forcing its rotation in the case of gp33. The TCR residue Y101α, which side chain is positioned between p1K and p4Y, forms a hydrogen bond with the H-2D^b^ residue E163, which also forms a hydrogen bond with p4Y (Fig. S7). Thus, the hydroxyl group of p4Y plays a key role in a net of hydrogen bond and van der Waals interactions formed with TCR residues N38a and Y101α as well as the H-2D^b^ residue E163. The Y4F mutation abrogates all these interactions, abolishing P14 recognition (Fig. S7). Furthermore, the Y4F mutation introduces high hydrophobicity within this key TCR/pMHC interface, composed mainly by polar residues. Altogether, this explains why P14 does not bind nor recognize the immune escape variant H-2D^b^/Y4F.

### The p3P modification facilitates TCR recognition

The three ternary TCR/MHC/peptide structures were compared with each corresponding TCR-unbound pMHC (Fig. 4, Fig. S8). The side chain of p4Y, essential for recognition by P14 [4, 33, 34], rotates down following P14 binding to both H-2D^b^/gp33 and H-2D^b^/V3P (Fig. 4A-B). A similar rotation was also observed for residue p4F in H-2D^b^/PF upon binding to P14 (Fig. 4C). The side chain of p6F in gp33 is also affected upon binding to P14 (Fig. 4A). Interestingly, the p3P modification resulted in a similar conformation for p6F in both H-2D^b^/V3P and H-2D^b^/PF prior to binding to P14 (Fig. 1D and Fig. 4). Furthermore, the side chain of residue p1K in H-2D^b^/gp33 also moves towards the N-terminal of the peptide binding cleft following P14 binding (Fig. 4A), taking an identical conformation as in both p3P-substituted peptides (Fig. 4D).

**Figure 4.**
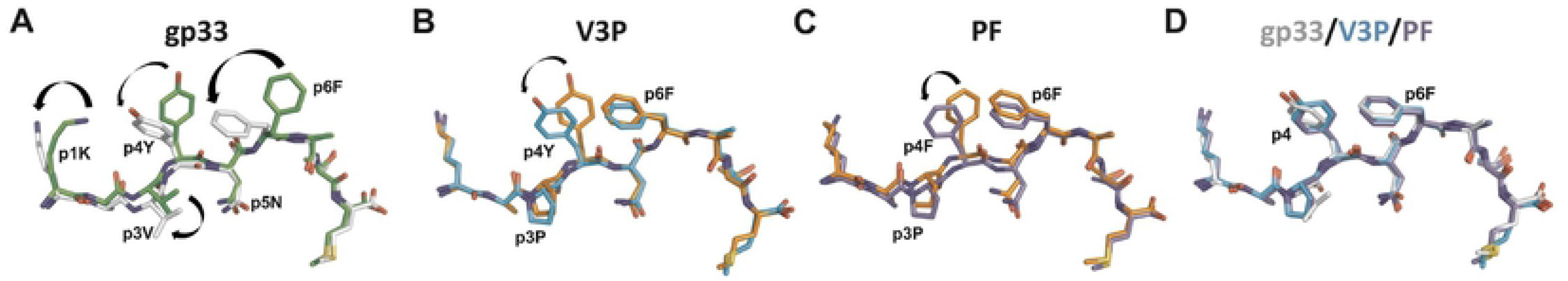
The p3P modification results in conformational changes of peptide residues p1K and p6F, predisposing pMHCs for optimal binding to P14. **A.** Comparison of gp33 before binding (in green) and after binding (in white) to P14 reveals major conformational changes in gp33 following binding to P14. These include a movement of the p2-p4 backbone of gp33 that is pushed down in the cleft combined with a 180 degrees rotation of the isopropyl moiety in residue p3V. Furthermore, the side chain of peptide residues p1K, P4Y and p6F all take new conformations following binding to P14. All movements are indicated by blue arrows. **B.** The introduction of p3P in V3P results in optimal positioning of the side chains of residues p1K and p6F prior to binding to P14 (in orange). The only observed conformational difference was taken by residue p4Y following V3P binding to P14 (in cyan). **C.** Similarly to V3P, the only conformational difference observed for PF before (in orange) and after (in violet) binding to P14 is at peptide residue p4Y. **D.** Peptides gp33 (in white), V3P (in cyan) and PF (in violet) take nearly identical conformations when bound to P14.

One of the most significant differences in H-2D^b^/gp33, before and after binding to P14, is a shift of the p2-p4 backbone of gp33 when bound to P14, towards the binding cleft of H-2D^b^. Following P14 docking, p3V in gp33 extends 1.2 Å deeper into the D-pocket of H-2D^b^, combined with a 180° rotation (Fig. 4A). In contrast to gp33, the p2-p4 section is more constrained in both V3P and PF, following TCR binding (Fig. 4B, 4C). However, it should be noted that the final conformations of all three peptides in the ternary complexes is nearly identical (Fig. 4D). In conclusion, residues 1 and 6 in p3P-APLs take the same conformations prior to TCR binding as found in the ternary complexes, potentially enabling a more favorable surface for P14 TCR binding.

The crystal structures of TCR-unbound and TCR-bound pMHCs also revealed that conformational differences in H-2D^b^ residues were observed only for residues R62, E163 and H155 (Fig. 5, S2, S9 and S10). The large movement of p6F in gp33 following binding to P14 induces the counter wise reorientation of the side chain of residue H155 towards the TCR (Fig. 5A). The redisposition of H155 and p6F in H-2D^b^/gp33 promotes the adequate positioning of the key TCR residue R97b, which runs longitudinally along the length of the N-terminal part of the peptide (Fig. S7). In contrast, residues p6F and H155 are already optimally positioned in both the TCR unbound and bound forms of the H-2D^b^/V3P and H-2D^b^/PF complexes (Fig. 5B–5C), most probably predisposing for optimized interactions with P14.

**Figure 5.**
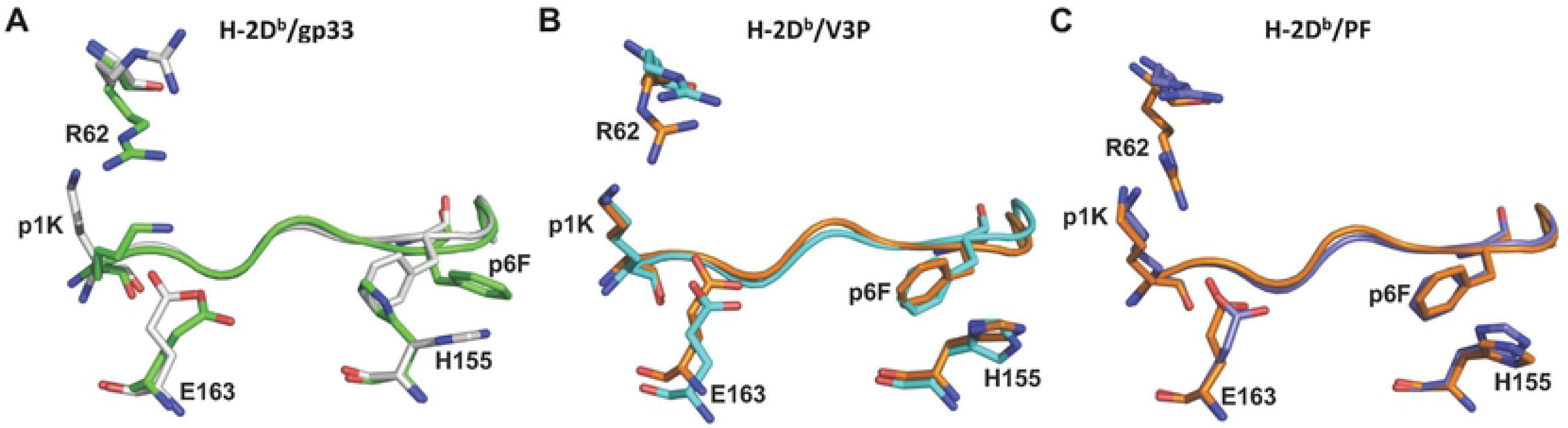
The p3P modification affects the conformations of peptide residues p1K and P6F, as well as H-2D^b^ residues R62, H155 and E163 facilitating TCR recognition. **A.** Comparison of H-2D^b^/gp33 before (in green) and after P14 binding (in white) reveals that the conformation of a very limited amount of pMHC residues is affected (shown as sticks). Following binding to P14, the side chain of peptide residue p1K moves towards the N-terminal part of the peptide binding cleft while the side chain of p6F rotates. As a consequence, conformational changes are observed only for heavy chain residues R62, H155 and E163. **B.** In contrast to gp33, the introduced p3P modification already positions most peptide and heavy chain residues in optimal conformations, limiting significantly the required movements following binding to P14. pMHC residues before and after binding to P14 are colored orange and cyan, respectively. **C.** Similarly to V3P, the p3P modification in PF results in optimal positioning of all key peptide and heavy chain residues prior to binding to P14. pMHC residues before and after binding to P14 are colored orange and violet, respectively.

Furthermore, p1K in gp33 also takes a different conformation upon binding to P14, bending backwards towards the H-2D^b^ residues R62 and E163, which conformations are affected (Fig. 4A, 5A). Here again, the side chain of p1K takes exactly this conformation in both V3P and PF already before TCR binding (Fig. 4, 5). Altogether, p1K, P6F and heavy chain residues R62, H155 and E163 have already adopted in the unbound V3P and PF complexes similar conformations to those observed in all three ternary structures (Fig. 4, 5). Thus, the p3P substitution potentially facilitates TCR recognition by positioning specific key peptide and MHC residues prior to the formation of the ternary complexes. This is well in line with our SPR and ITC results, which indicate that the energy required for P14 recognition of V3P is reduced compared to gp33 (Table 1, Fig. S1).

## Discussion

Subsets of peptide analogs have been used by others and us to both break T cell tolerance and enhance T cell responses to tumors [16, 23, 35]. Heteroclitic subdominant viral T cell determinants were also used to enhance both pMHC stability and T cell avidity towards the mouse hepatitis virus-specific subdominant S598 determinant [22, 36]. Most, if not all studies performed in other laboratories have focused their efforts on introducing peptide mutations that would significantly increase the stability of pMHCs with as little alteration as possible of peptide conformation. Here, instead of mutating a key anchor position, we targeted interactions between peptide position 3 and the MHC residue Y159, conserved among most known mouse and human alleles. Indeed, besides H-2D^b^, we have previously demonstrated that the p3P modification enhances significantly the stability of H-2K^b^ in complex with different TAAs [23]. Thus, the p3P modification could potentially enhance stabilization of other MHC-I alleles that comprise the heavy chain residue Y159, leading to enhanced TCR recognition.

Here, we addressed if we could increase CD8^+^ T cell avidity and restore recognition of the viral escape variant Y4F that binds to H-2D^b^ with the same high affinity as gp33 [37]. The TCR P14 is specific for H-2D^b^/gp33 and it has been previously demonstrated that P14 recognition is abolished by the Y4F mutation [2, 3]. Comparison of the crystal structures of H-2D^b^/gp33 and H-2D^b^/Y4F demonstrated that the only structural difference between these two pMHCs was the removal of the hydroxyl tip from the peptide residue p4 [2]. We demonstrate that the p3P modification in PF overcomes the restrictions imposed by the Y4F mutation, reestablishing P14 recognition of this structural mimic of Y4F. Furthermore, we show that it is fully possible to restore endogenous CD8^+^ T cell recognition of Y4F following vaccination with PF. Possibly, the higher avidity of subsets of the endogenous T cell population for H-2D^b^/PF pushes them over a certain threshold of activation, and the molecular similarities between H-2D^b^/PF and H-2D^b^/Y4F allow for crossreactivity, resulting in significant cytokine secretion towards Y4F. However, *in vitro* re-stimulation with gp33 of endogenous CD8^+^ T cells isolated from Flu(PF)-vaccinated mice did not induce any significant secretion of cytokines, although these endogenous CD8^+^ T cells recognized both PF and gp33-loaded MHC tetramers. Martin *et al* have previously provided evidence for selective activation of different effector functions in CD8^+^ T cells by APLs. More specifically, the results of their study show that the H-2Db/Y4F-specific T cell clone YF.F3 killed efficiently targets presenting gp33 but did not produce high amounts IFNγ against gp33 [32]. This is well in line with the results presented in this study. Altogether, this suggests to us that vaccination with a cocktail of epitopes could provide wider protection against both immunodominant and immune escape targets.

So how does it possibly work at the molecular level? The rigidification of p3P-modified peptides could enhance TCR recognition by decreasing entropic costs. Indeed, we have previously demonstrated in TAA models that peptide rigidification enhanced considerably TCR recognition [26, 27]. Overall the effects of proline replacement on protein stability and function are well established for a large ensemble of proteins [38], revealing that protein-protein interactions often occur through regions enriched with proline residues [39]. Proline substitutions increase overall protein stability as well as the stability of specific protein regions [40]. Indeed, proline replacement of specific residues in TCR CDR loops can increase significantly recognition of antigens [41]. The importance of the interaction of peptide residue p3P with residue Y159, conserved amongst most known MHC-I alleles, has been previously described [27], revealing that p3P reduces significantly the flexibility of the pMHC complex, thus decreasing unfavorable entropic change upon complex formation. Such reduced entropic penalties for TCR recognition following p3P mutation were confirmed here by ITC measurements, which indicated reduced unfavorable entropic contribution for recognition of H-2D^b^/V3P by P14 compared to H-2D^b^/gp33. The importance of the reduction of peptide conformation heterogeneity for enhanced TCR has been described, using a combination of crystal structure and molecular dynamic studies [42]. A peptide that must move to optimize the interactions with the bound TCR will increase the entropic cost for binding, resulting in slower binding, lower affinity and less efficient recognition [43]. Consequently, although many TCRs bind with unfavorable entropy changes [37, 44], reduction of conformational heterogeneity coupled with rigidification of the peptides may lead to enhanced T cell recognition. In this study, the p3P mutation reduces motion and therefore enhances T cell recognition by increasing T cell association rate and decreasing entropic costs for binding.

Although X-ray structural studies of proteins provide accurate snapshots of protein complexes, crystal structures provide relatively little information about the dynamic bases underlying protein-protein interactions. The dynamic motions of both pMHC and TCR impact on recognition by T cells, clearly influencing function and recognition [42]. Here, we compared the crystal structures of each studied pMHC complex before and after P14 TCR binding (besides the P14/H-2D^b^/Y4F complex that could not be obtained since P14 does not bind to this pMHC). Peptides tune the motions of MHC heavy chains and reduced motions may lead to enhanced recognition. Besides the peptide rigidification imposed by the p3P modification, comparison of a structural snapshot for each ternary structure with each TCR-unbound pMHC variant indicated an additional structural reason for the increased TCR recognition of p3P-modified epitopes. In all cases, conformational differences were observed in peptide residues p1K and p6F in PF and V3P, compared to Y4F and gp33, before TCR binding. In all p3P cases, the side chains of peptide residues p1K and p6F took the same conformation, as observed in the ternary structures, prior to TCR binding. In line with this, others [33, 45] and we [37] have previously demonstrated the importance of residue p1K for recognition by the TCR P14. The crystal structure of the semi-agonist Y4A (KAVANFATM) also revealed a similar conformation for both p1K and p6F prior to binding to P14 TCR [37]. Furthermore, the conformation of the MHC “TCR footprint” heavy chain residues R62, H155 and E163 [46, 47] was also affected following p3P substitution, possibly due to the movements of p1K and p6F. Altogether, prior to TCR landing, the p3P modification alters the conformation of residues both in the peptide and the MHC heavy chain similar to conformations taken upon binding to the TCR, thus predisposing the pMHC for facilitated TCR recognition.

The results presented within this study indicate in our opinion that i) docking of P14 to p3P-modified peptides is facilitated since the conformations of key residues in both peptide and heavy chain are already optimal prior to TCR binding (ready-to-go conformation); ii) consequently, the energetic costs for TCR recognition should be reduced since there is no need for any major movement in the rigidified epitope besides the conformational change for residue p4Y. As vaccination with PF restored endogenous T cell recognition of Y4F, the p3P modification could thus represent a novel way to increase the immunogenicity of a large array of H-2D^b^-restricted epitopes as well as possibly viral epitopes restricted by other MHC alleles. We thus describe here a successful approach to restore recognition of viral escape peptide that can be easily coupled to already existing vaccination protocols, including vaccination with full-length proteins as well as *e.g*. modified mRNA vaccines, by introducing the p3P modification in a selection of viral epitopes.

## Materials and Methods

### Cell lines and mice

H-2D^b+^/H-2K^b+^ RMA cells, kindly provided by Prof. Klas Kärre, were used as target cells in the functional assays described below. Pathogen-free wild-type (WT) C57BL/6 (B6) and RAG1/2-deficient (RAG1/2^-/-^) P14-transgenic mice were bred and maintained within the facilities of the MTC department, Karolinska Institute. Vα2^+^ T cells from P14 mice were used as effector cells for *in vitro* experiments. P14 mice were used for *in vivo* T-cell stimulation assays.

### Peptides and antibodies

Peptides gp33, Y4F, V3P and PF as well as control peptides NP_366_ (ASNENMETM) and P18-I10 (RGPGRAFVTI) were purchased from GenScript (Piscataway, NJ, USA). Antibodies 53-6.7 (anti-CD8α), 53-5.8 (anti-CD8β), XMG1.2 (anti-IFN-γ), MP6-XT22 (anti-TNF), 145-2C11 (anti-CD3ε), 1D4B (anti-CD107a), BP-1 (anti-Ly5.1/CD249), IM7 (anti-CD44) and B20.1 (anti-TCR Vα2) were purchased from BD Biosciences (San Diego, CA, USA). Antibodies GK1.5 (anti-CD4) and H57-597 (anti-TCR Cβ) were purchased from Abcam (Cambridge, UK) and eBioscience (San Diego, CA, USA), respectively.

### Preparation, refolding and crystallization of TCR/pMHC complexes

Refolding of all pMHCs was conducted as previously described [48]. P14 was produced and refolded by dilution and thereafter-purified using ion exchange and size exclusion chromatography. Crystals for H-2D^b^/V3P and H-2D^b^/PF were obtained by hanging drop vapor diffusion in 1.6-1.8 M ammonium sulfate, 0.1 M Tris HCl pH 7.0-9.0. Crystals for P14/H-2D^b^/gp33, P14/H-2D^b^/V3P and P14/H-2D^b^/PF were obtained by hanging drop vapor diffusion in 19% PEG 6000, 0.1 M Tris HCl pH 8.0.

### Data collection, processing and structure determination

Data collection was performed at beam lines ID14-2 and ID23-2 at ESRF (Grenoble, France). Diffraction data were processed and scaled using MOSFLM 7.0.3 and SCALA [49]. Crystal structures were determined by molecular replacement using PHASER [50]. The crystal structure of H-2D^b^/gp33 (PDB ID: 1S7U) [2], with omitted peptide, was used as search model for H-2D^b^/V3P and H-2D^b^/PF. P14/H-2D^b^/gp33, P14/H-2D^b^/V3P and P14/H-2D^b^/PF were determined using 3PQY [51]. In all cases, poorer electron density was displayed for the TCR Cα domain, probably due to high flexibility, as previously observed [52]. Random 5% reflections were used for monitoring refinement by R_free_ cross-validation [53]. The model was rebuilt in Coot where necessary. The stereochemistry of the final models was verified using PROCHECK [54] or Coot [55].

### Circular dichroism (CD) analysis

Measurements were performed in 20mM K_2_HPO_4_/KH_2_PO_4_ (pH 7.5) using 0.15-0.3 mg/ml protein concentrations. Melting temperatures (Tm) were derived from changes in ellipticity at 218 nm as previously described [37]. Curves and T_m_ values are an average of at least three measurements from at least two independent refolding assays per pMHC. Spectra were analyzed using GraphPad Prism 5 (La Jolla, USA).

### Surface Plasmon Resonance (SPR) binding affinity analysis

All measurements were performed on BIAcore 2000 (GE Healthcare, USA) at 25°C. Soluble P14 (20 μg/ml) was non-covalently coupled to the anti-C_β_ antibody H57-597. 8000 RUs of H57-597 were coupled to a CM5-chip, resulting in 3000RUs immobilized P14. A control surface without antibody was used as reference. Concentration series of pMHCs were injected over the chip. The surface was regenerated with 40 μl 0.1 M Glycine-HCl, 500 mM NaCl, pH 2.5. Unspecific binding was corrected for by subtracting responses from reference flow cells. Data were analyzed with BIAevaluation 2000 (BIAcore AB, Uppsala, Sweden). K_D_-values were obtained from steady-state fitting of equilibrium binding curves from at least two independent measurements.

### Isothermal titration calorimetry (ITC)

Measurements were performed on a MicroCal iTC 200 (GE Healthcare, USA) at 25°C. 40 μl H-2D^b^/V3P (125μM) or H-2D^b^/gp33 (150 μM) in 10 mM Hepes, 150 mM NaCl, pH 7.4 were titrated into 300 μl of P14 (12.5-15 μM) in 10 injections under 1000 rpm stirring rate. Data analysis was performed using Origin, fitted to a non-linear curve in an iterative process. The reported constants are an average of two independent experiments.

### TCR down-regulation assays

P14-splenocytes were mixed with peptide-pulsed RMA cells at 10:1 effector:target (E:T) ratio. Cells were co-incubated at 37°C for 4 h and stained with anti-CD8β and -TCR Vα2 antibodies. Flow cytometry was performed using FACSCalibur (BD Biosciences) and changes in mean fluorescence intensity (MFI) of the Vα2 staining were used to estimate TCR down-regulation. Data was analyzed using Flowjo (Tree Star, Inc., Ashland, OR, USA).

### *In vivo* stimulation of P14 T cells and Cr^51^ release cytotoxicity assays

P14 TCR-transgenic mice were injected subcutaneously (SC) with 100 μg gp33 in PBS combined with 12.5 ng phosphorothioate-modified CpG-ODN 1668 (Invivogene, Sweden). 20 mg Aldara cream was applied at site of injection (5% imiquimod, Meda AB, Sweden). Animals were sacrificed 7 days later and spleens were recovered. Target RMA cells, labeled with Cr^51^, were pulsed with indicated peptide concentrations for 1 h at 37°C and subsequently mixed with *in vivo*-stimulated negatively selected (MACS CD8^+^ T cell isolation kit, Miltenyl Biotec, Germany) P14 CD8^+^ T cells at 3:1 E:T ratio followed by a standard 4h Cr^51^-release assay. Radioactivity was measured on a γ-counter (Wallac, Uppsala, Sweden). Percentage of specific lysis was calculated as [Cr^51^ release in test well – spontaneous Cr^51^ release] / [maximum Cr^51^ release – spontaneous Cr^51^ release] x 100.

### CD107a degranulation, intracellular IFNγ and TNF production

CD8^+^ T cells isolated from spleens of naïve or *in vivo*-stimulated P14 transgenic mice were cocultured for 5 h with 10^-6^ M or 10^-8^ M peptide-pulsed RMA cells in the presence of anti-CD107a antibody for degranulation assays. GolgiStop (BD Biosciences) was added after 1 h co-incubation. 4 h later, cells were stained with anti-CD8α and -CD3ε antibodies. For intracellular cytokine staining assays, cells were fixed and permeabilized using the Cytofix/Cytoperm kit (BD Biosciences) according to instructions. Cells were thereafter stained for IFNγ and TNF expression. FACS sampling was performed on CyAn (Dako, Glostrup, Denmark) and analyzed with FlowJo.

### MHC-I tetramer production

H-2D^b^ molecules with a biotinylation tag were refolded with peptides and mβ_2_m in the presence of protease inhibitors and purified as previously described [56]. Each obtained monomeric H-2D^b^/peptide complex (0.5 mg/ml) was tetramerized at a 4:1 ratio with streptavidin-PE (BD Biosciences).

### Identification of P14 CD8^+^ T cell responses upon LCMV vaccination

10^4^ P14 T cells (CD44^low^, Ly5.1^+^), isolated from spleens of P14 transgenic mice, were adoptively transferred intravenously (in PBS) into C57Bl/6 mice three days prior to intraperitoneal infection with 1×10^6^ PFU of LCMV (Clone 13). Spleens were harvested on day 7 post infection. CD8^+^ T cells were enriched by B cell panning and red blood cell lysis and stimulated with IL-2 (25 units/ml), Brefeldin A (5 μg/ml) (BD Biosciences) and 10^-6^ M peptide (gp33, Y4F or PF or no peptide) in complete RPMI for 5 h at 37°C, 5% CO_2_. Washed cells were surface stained with anti-CD8, -CD44 and -Ly5.1, fixed and permeabilized using BD cytofix/cytoperm kit (25 min at 4°C). Intracellular staining of IFNγ, TNF and IL-2 (at 1:200) was performed for 30 min at 4°C. Endogenous T cells were distinguished by congenic marker Ly.5.2 from Ly.5.1^+^ P14 T cells. Cells were resuspended in FACS buffer after enrichment and stained at 1:400 for 1hr at RT with gp33, Y4F or PF tetramers. Washed cells were surface stained for CD8, Ly5.1, CD107a and CD44 for 30 min at 4°C. Data was collected using LSR Fortessa (BD Biosciences) and analyzed with Flowjo.

### Cloning of Plasmids

pHW2000 vectors containing the 8 genes (PB2, PB1, PA, HA, NP, NA, M and NS), where NA and HA are derived from HKx31 (H3N2), and the internal genes from A/PR8/34 (PR8, H1N1), were constructed by reverse transcriptase-PCR (RT-PCR) amplification of the viral RNA. The peptides Y4F and PF were introduced into the Influenza A virus by inserting/replacing a region in the stalk of Neuraminidase (NA) using the cloning system as described.

### Viruses and Cell Culture

Reverse genetics, generation of modified Influenza: Briefly, 1 ug of each plasmid (NP, NS2, PB2, M, PA, PB1, HA and NA) was mixed with 16ug of lipofectomine and OptiMEM and added to a mix of co-cultured MDCK/293T cells, in the presence of TPCK-trypsin. The transfection was allowed to proceed for 48-72h in 5% CO_2_ at 37°C. The virus was thereafter propagated in chicken eggs for 2 days at 35° [57].

### RNA Isolation and RT-PCR

Viral RNA was isolated from virus particles with RNeasy-Kit (Qiagen, Valencia, CA). Access RT-PCR kit (Promega) was used for characterization of recombinant influenza viruses.

### Identification of T cell responses upon Influenza vaccination

Naive C57Bl6 mice were adoptively transferred with 10^4^ P14 T cells one day prior to infection. with 1 x 10^4^ PFU or i.p. with 1.5 x10^7^ PFU of influenza A virus following anesthesia with isofluorane, then used for analysis of primary immunity at day 10 post infection. Kinetics, magnitude and phenotype of primary virus-specific CD8^+^ T cell responses were measured by flow cytometry. gp33- and APL-specific CD8^+^ T cell populations were characterized using H2D^b^/gp33, Y4F and PF tetramers. Splenocytes were incubated with tetramers for 60 min at room temperature. Washed cells were stained for CD8^+^ and CD44 for 30 min at 4°C. Intracellular IFNγ and TNF staining (1:200) was performed for 30 min at 4°C. Data was collected using LSR Fortessa (BD Biosciences) and analyzed with Flowjo.

### Statistical analysis

Data were routinely shown as mean ± SD. Unless stated otherwise, statistical significance was determined by the Student’s t test or analysis of variance (ANOVA) using GraphPad Prism 7.0. *P < 0.05; **P < 0.01; ***P < 0.001; ****P < 0.0001.

### Ethics statement

All experimental animal procedures were performed under Swedish national guidelines (N413/09) and following approval from the University of Melbourne animal ethics experimentation committee (ethics number 1312890.3).

## Abbreviations

APL: altered peptide ligand
BAL: bronchoalveolar lavage
CD: circular dichroism
CTL: cytotoxic T-lymphocyte
ITC: isothermal titration calorimetry
LCMV: lymphocytic choriomeningitis virus
MHC-I: major histocompatibility complex class I
pMHC: peptide/MHC complex
SPR: surface plasmon resonance
TCR: T-cell receptor
TEIPP: T cell epitope associated with impaired peptide processing

